# Risk assessment during nest defense against three simulated predators by female northern house wrens (*Troglodytes aedon*)

**DOI:** 10.1101/2023.02.01.526454

**Authors:** Ross C. Eggleston, Josephina H. Fornara, Kyle M. Davis, Jess Dong, Dustin G. Reichard

## Abstract

Offspring predation is one of the greatest obstacles to an organism’s reproductive success, but parents vary in the strength of their response to potential predators. One explanation for this variable investment is that defending current offspring has the potential to lower future reproductive success if the predator is also capable of injuring or killing the parent. Northern house wrens (*Troglodytes aedon*) are cavity-nesting songbirds that defend against multiple species of nest predators including small mammals, birds of prey, and snakes. Here, we used three different predator decoys: two nest predators - an eastern chipmunk (*Tamias striatus*) and an eastern ratsnake (*Pantherophis alleghaniensis*) - as well as a predator of both offspring and adults - a juvenile Cooper’s hawk (*Accipiter cooperi*) - to elicit nest defense and test whether females use risk assessment to modulate their antipredator behavior. We found that antipredator behaviors were not significantly different between the two nest predators, which posed a high risk to the nestlings, but lower risk to the parents as neither species frequently captures adult wrens outside the nest box. However, female wrens never dove at or attacked the Cooper’s hawk while they frequently attacked both the snake and chipmunk decoys. Neighboring house wrens from adjacent territories were also less likely to respond to the hawk, but more heterospecifics mobbed the hawk than the snake decoy. Collectively, these results show that risk assessment and the strength of the antipredator response varies substantially both within and among species. Female house wrens exhibit plasticity in their nest defense behavior, and they respond to different types of predators in a way that could maximize lifetime fitness while risking the loss of their current offspring.

## Introduction

Parental investment can be defined as any behaviors that increase an offspring’s chance of survival while simultaneously decreasing the parent’s ability to invest in other offspring both currently and in the future (Trivers 1974; Montgomerie and Weatherhead 1988; Clutton-Brock 1991). This tradeoff between current and future reproduction creates a parent-offspring conflict (*sensu* Trivers 1974) because the optimal parenting strategy, one that maximizes the parent’s lifetime fitness, often requires submaximal investment in the current offspring, which can decrease their survival and fitness. The amount of parental investment varies widely both within and among species (Fontaine and Martin 2006; Dulac et al. 2014), and this variation can be further explained by differences in life history strategies (Ricklefs 1977; Montgomerie and Weatherhead 1988; Clutton-Brock 1991; Ghalambor and Martin 2001). For example, both adult mortality rates and offspring number and age should correlate positively with parental investment because species with shorter lifespans and larger current investments will have fewer reproductive opportunities in the future (Caro 2005).

Optimally modulating parental investment is particularly critical in the context of antipredator behavior, as parents that vigorously defend their young risk completely eliminating their future reproduction if they are mortally injured or killed. On the other hand, parents that limit their antipredator behavior are more likely to lose their current offspring as predation is one of the most common causes of reproductive failure across species (Ricklefs 1977; Caro 2005; Martin and Briskie 2009; Ibáñez-Álamo et al. 2015). The most successful parents should be those that exhibit plastic antipredator behavior that maximizes their lifetime reproductive success, and selection should favor individuals with accurate risk assessment during encounters with predators near their offspring (Montgomerie and Weatherhead 1988).

An abundance of evidence supports the hypothesis that animals from across the taxonomic spectrum use risk assessment to adjust their antipredator behavior relative to the perceived threat level (reviewed in Caro 2005; Lima 2009). Species with altricial young, such as passerine songbirds, are particularly vulnerable to predation at the nest because their offspring have limited capabilities for self-defense or escape without parental assistance (e.g., Suzuki 2011). Consequently, parents respond to different predation pressures with both proactive and reactive behavior. When the perceived risk of predation is high, parents can proactively reduce egg or clutch sizes (Fontaine and Martin 2006; Morosinotto et al. 2010; Travers et al. 2010) or limit visits to the nest, including offspring provisioning (Fontaine and Martin 2006; Ghalambor et al. 2013; Dudeck et al. 2018), which minimizes their potential energetic losses and draws less attention to the nest site. Alternatively, parents may simply avoid nesting in locations with high predator abundance or even forgo reproduction entirely (Spaans et al. 1998; Morosinotto et al. 2010; Szymkowiak and Thomson 2019).

When a predator appears at the nest site, parents can react in ways that might divert the predator’s attention toward the parents rather than the offspring such as alarm calling or visual distraction displays (e.g. broken-wing display; de Framond et al. 2022). Alarm calling also has the potential to initiate a community-level response by attracting nearby conspecifics and heterospecifics to collectively harass (i.e., “mob”) the predator, which may increase the likelihood of expelling the predator from the area (Carlson and Griesser 2022). Although alarm calls and visual displays increase risk to the parents, they are often less costly than more aggressive defense behaviors such as diving at or hitting the predator in an attempt to physically drive it away from the offspring. Studies from multiple species have shown that parents exhibit plasticity in their antipredator responses and adjust their behavior according to factors such as the distance between the predator and the offspring (Colombelli-Négrel et al. 2010; Fernández and Carro 2022) and the relative risks posed to both adults and offspring by different types of predators (Gottfried 1979; Winkler 1992; Mahr et al. 2015; Duré Ruiz et al. 2018; Teunissen et al. 2020). Similarly, the willingness of nearby conspecifics and heterospecifics to respond to alarm calls with mobbing behavior varies according to the intensity of the calls (Templeton et al. 2005, Fernández et al. 2023) and the risks posed by the predator (Forsman and Mönkkönen 2001, Scott and Robinson 2023). In this study, we tested for evidence of risk assessment by female northern house wrens (*Troglodytes aedon*) during simulated interactions with predators at their nest that posed different threats to adults and nestlings.

Previous research in both northern and southern house wrens (Troglodytes musculus) has found evidence for risk assessment and plasticity in parental behavior in response to predation pressure. For example, wren parents produced more alarm calls when an owl model was placed closer to their nest (Fernández and Carro 2022), consistent with an ability to perceive levels of risk. After interacting with a simulated nest predator, wren parents have also been shown to reduce their offspring provisioning (Freed 1981; Ghalambor et al. 2013) and take longer to re-enter the nest box when compared to control observations (Corral et al. 2013; Fernandez and Llambías 2013; Fernández et al. 2015; Duré Ruiz et al. 2018). When presented with different simulated species of raptors of varying threat levels, male and female southern house wrens took longer to resume normal parental behavior after encountering a predator of both adults and nestlings than a predator of nestlings only (Duré Ruiz et al. 2018). This result suggests that risk assessment may shape parental behavior in adult wrens by shifting investment away from offspring and towards self-preservation when perceived threat levels are high.

Here, we expand on these observations by testing whether wren parents assess risks posed by both avian and non-avian predators and adjust the strength of their nest defense relative to risks associated with each type of predator (see below). Over two breeding seasons, we placed three simulated predators of varying risks on top of house wren nest boxes during the early nestling stage. One predator, a rubber eastern ratsnake decoy (*Pantherophis alleghaniensis*), was presented in both years, while the second predator differed between years and consisted of a rubber eastern chipmunk decoy (*Tamias striatus*) in year one and a taxidermied juvenile Cooper’s Hawk (*Accipiter cooperii*) in year two. We assumed that the ratsnake and chipmunk decoys posed a high risk to nestlings, but a low-to-moderate risk to adult wrens, which are unlikely to be captured by either predator outside of the nest box (Reidy et al., 2009; Johnson, 2020). In contrast, the hawk posed a high risk to adults, but a low-to-moderate risk to nestlings, which are still vulnerable to raptors and other avian predators despite the protection offered by an enclosed nest box (Milsap et al. 2013; Johnson 2020). If the hypothesis that house wrens use risk assessment was supported, we predicted that female wrens would respond equally aggressively to the ratsnake and chipmunk decoys, which are similar in threat level to both nestlings and adults. Female responses to the hawk were predicted to be less aggressive overall due to the higher threat that the hawk posed to adults and lower threat posed to nestlings. Finally, we examined the community-level response to predators of varying risk by quantifying the number of conspecific and heterospecific individuals that responded to each predator treatment.

## Methods

### Study System and Sites

Northern house wrens (*Troglodytes aedon*) are socially monogamous, cavity-nesting songbirds that readily use artificial nest boxes to raise clutches of up to 8 offspring (Johnson 2020). The guild of house wren nestling predators is diverse and includes mammals ranging in size from mice (e.g., *Peromyscus* spp.) to bears (*Ursus spp.*), passerines (e.g., *Cyanocitta* spp.), woodpeckers (e.g., *Melanerpes* spp.), raptors, and snakes (Johnson 2020). At our study sites the most common house wren nest predators are small mammals and snakes, but multiple raptor species also frequent our sites and pose risks to adults and offspring (DGR, personal observation).

We monitored house wren breeding success at 180 nest boxes divided between six locations in Delaware and Union Counties, Ohio, USA (Table S1, Supplementary Materials) during the 2020 and 2021 breeding seasons. The nest boxes varied slightly in dimensions due to landowner preferences, but the average internal dimensions were 110 x 115 x 195 mm and the entrance hole size was 38 mm. Each nest box was mounted on a conduit pipe approximately 1.75 m above the ground, and the pipe was covered in axle grease to limit access by predators. The majority of the boxes were located in edge habitat near the transition from grass or field to forest, but a small subset was located in suburban backyards (*N*=7).

### Banding of Adults and Nestlings

We captured male house wrens using mist nets and a brief playback of conspecific song. Each male was banded with a USFWS aluminum leg band and a unique combination of 2-3 additional colored bands for identification from a distance. We also collected morphological measurements and a small blood sample for an unrelated study. Banding only males allowed us to differentiate between the sexes during our behavioral trials (see below), and because most females were never captured, it also minimized the likelihood that the capture process disrupted the female’s antipredator response. Any females that were captured accidentally (Experiment 1, *N* = 6 of 38 females; Experiment 2, *N* = 10 of 38 females) were rapidly banded and immediately released within a few minutes of capture to limit any potential effects of handling on subsequent behavior. All birds were captured and banded at least 24 hours before any behavioral trials, and most females that were captured were handled more than one week before their first trial (Experiment 1, *N* = 4/6; Experiment 2, *N* = 5/10). We assumed that the small number of females captured, the limited handling time, and the delay after any female was captured before conducting behavioral trials minimized any effects of unintentional handling on female responses.

We also monitored the number of eggs laid, hatching success, and offspring development for every nest box. Nest checks occurred every 3-4 days, and any checks on the day of a behavioral trial were conducted after the trial concluded to minimize any effect on behavior.

We counted the number of nestlings at the end of each behavioral trial (see below), and banded nestlings when they were 8 days old. Each nestling received a single USFWS aluminum band, and we collected morphological measurements and a small blood sample. After banding, we passively monitored each nest box until all visible and acoustic activity ceased. Unless the contents of the nest suggested otherwise (i.e., presence of dead nestlings and/or their parts), we assumed that empty nests successfully fledged.

### Experiment 1: Chipmunk v. Snake Decoy

In 2020, we conducted behavioral trials between June 9 and July 24 at 47 nest boxes containing nestlings between the ages of 3-7 days old. A smaller age range was impossible due to the distances between our sites and other field-related challenges (e.g., weather). Two life-like rubber predator decoys, an eastern chipmunk (*Tamias striatus*; Safari Ltd.) and an eastern ratsnake (*Pantherophis alleghaniensis*; Flybanboo), were used to elicit nest defense behavior. We chose these predators because both species are common at our study sites and had been previously observed either depredating nests (ratsnake) or being actively attacked by house wrens near nests (both predators; personal observations). This comparison allowed us to empirically test the prediction that wrens should respond equally to predators of similar risk despite the fact that the two predators are morphologically and ecologically distinct.

To initiate a trial, we waited for each female to enter or land on the nest box before flushing them away during setup. One predator was placed on top of the nest box, and then on the following day the other predator was presented in a counterbalanced order to control for order effects. Although these stimuli are terrestrial predators, both are capable of climbing nearby vegetation and approaching the nest box from above, making their appearance on top of the box biologically relevant.

All trials occurred between 7:00-13:00, but we were unable to conduct trials exactly 24 hours apart due to the distant locations of our field sites and unpredictability in how rapidly females would return to their boxes. Trials were recorded with a video camera mounted on a tripod placed roughly 2-3 meters away from the nest box to later confirm behaviors and differentiate between the male and female wren in cases of uncertainty. We positioned the camera in tall grass or other vegetation to minimize its disruption to the focal birds, and observers maintained a distance of roughly 15 m away from the nest box. Each decoy was attached to a fishing line, and at the completion of the trial we used a fishing rod to rapidly reel the decoy away from the box. This approach eliminated the need for the observer to approach the box and remove the decoy. We allowed the video camera to continue to record for an additional 20 minutes to measure the amount of time it took for the female to re-enter the nest box after removal of the decoy (*N*=36). Post-trial data for some of the females were lost due to technical issues or user error.

We quantified the nest defense behavior of female house wrens for a total of seven minutes after the decoy was placed on the nest box. We counted the number of hits, defined as any contact the female made with the decoy, and the number of flyovers, defined as flights by the female over the decoy. We also measured the amount of time females spent within five meters of the nest box, the amount of time that females spent physically on or in the nest box, and the female’s closest approach (m) to the decoy. The five-meter threshold was chosen because all boxes had potential perches within five meters of the nest box despite variation in vegetative cover among our sites. Any female that hit the decoy was given a closest approach value of zero meters. We also noted whether or not the female produced alarm calls, and whether the female’s mate was present during the trial. Neighboring conspecifics and heterospecifics sometimes responded to the predator decoys by approaching, alarm calling, and in rare cases flying over and hitting the decoy. During each trial, we noted the number of individuals of each species that approached within 5 meters of the decoy. To avoid counting incidental birds that were not responding to the predator, we only counted individuals that either oriented towards the decoy or produced alarm calls.

Behavioral data from both predator trials for an individual female were excluded from our final analysis if either of the decoys were knocked off the box during either trial (*N*=3 of 47 total trials; 6.4%). However, if the chipmunk decoy was knocked over but remained on the box, the trials were not excluded (*N*=5 of 47 trials; 10.6%). We also excluded trials if the female did not return to the box within seven minutes of initiating the trial (*N*=6 of 47 trials, 12.8%). In response to this limitation, we adjusted our protocol early in the season so that we removed the decoy and reset the trial if the female did not return within 4 minutes after the trial was initiated. Once the female returned to the box, we flushed her away, placed the predator decoy, and started the seven-minute observation again. In summary, nine trials (of 47 trials; 19.1%) were excluded and our final sample size was 38 paired behavioral trials.

### Experiment 2: Hawk v. Snake Decoy

In 2021 between June 6 and July 30, we replicated this experiment in the same house wren population, but we replaced the chipmunk decoy with a taxidermied, juvenile Cooper’s hawk (*Accipiter cooperii*), which is commonly observed at our study sites and elicits alarm calls from passerines and small mammals (DGR, personal observation). Although Cooper’s hawks primarily forage on small or medium-sized adult songbirds and mammals, they have been frequently observed capturing nestlings from both cavity nests in bird boxes and open-cup nests in the forest canopy (Milsap et al. 2013, Rosenfield et al. 2020). Thus, the comparison of the response to the Cooper’s hawk versus the ratsnake allowed us to test the prediction that wrens should respond less aggressively to the higher risk predator of adults (hawk). The taxidermied hawk was mounted with wire onto a wooden plank, and two large binder clips were attached to the bottom of the plank with jewelry wire. This design allowed the decoy to be quickly attached and removed from the roof of the nest box while eliminating the possibility that it could be knocked off the box.

We conducted behavioral trials at 41 nest boxes with nestlings aged 2-7 days. A different observer (JHF) scored the trials in 2021, but she was trained by the observer (RCE) from 2020 to ensure consistency between years. All of our sampling methods were replicated from experiment 1, but we were unable to continue to use the fishing rod and line to rapidly remove the decoys. As a result, one of the observers quickly ran to the box and removed the decoy at the end of each seven-minute trial. The video camera continued to record for an additional 20 minutes (*N*=38), but the approach of the observer may have affected the return to normal behavior. The snake decoy was knocked off the nest box during three trials (of 41 total; 7.3%), and we excluded both the snake and hawk trials from those subjects in our final analysis for a total sample of 38 paired behavioral trials.

### Note on Experimental Design

Our decision to split this study into two separate experiments rather than one experiment including all three predator stimuli was driven by uncertainty stemming from the global Covid- 19 pandemic. This event shutdown numerous research labs in 2020 and disruptions continued into 2021. The potential for an unpredictable and abbreviated 2020 field season led us to err on the side of a less complicated protocol to maximize our sample size. We used the STRANGE framework to identify and address potential issues in sampling methodology in order to help prevent misinterpretation of results due to sampling biases (see Supp. Materials; Webster and Rutz 2020).

An additional limitation of this design was that we did not include a non-predatory control in either experiment (e.g., a heterospecific passerine that would pose no threat to a house wren nest). Thus, the behaviors we observed could be typical of a generalized nest defense response rather than a specifically “antipredator” response. We think this interpretation is unlikely for a few reasons. First, in a subsequent experiment, we compared the response of female wrens in our population to a novel object (car dice) placed on their nest box versus the ratsnake decoy. Female wrens responded significantly more aggressively to the snake and were significantly less likely to produce alarm calls when the novel object was present (DGR, unpublished data). This result suggests that female wrens in our population do not react to predatory and non-predatory stimuli with similar behaviors. Second, similar nest defense experiments in southern house wrens, a closely-related species, have shown that both male and female wrens respond less aggressively to non-predatory control birds at their nest (Fasanella and Fernández 2009, Duré Ruiz et al. 2017). Finally, a study in a different population of northern house wrens found that parents decreased their provisioning more in response to a simulated nest predator than a non-predatory control bird (Ghalambor et al. 2013). Although this study did not measure nest defense, it is consistent with wrens behaving differently around predators relative to non-predators near the nest.

### Statistical analysis

For each experiment, we generated either linear (LMM) or generalized linear (GLMM) mixed-effects models in R version 4.4.2 (R Core Team 2024; packages: “lme4,” “car,” “lmerTest”) to test whether female behaviors including flyovers, hits, closest approach to the decoy, time spent within five meters of the decoy, the production of alarm calls, and latency to re-enter the nest box after removal of the predator differed between predator types.

Measurements of time spent on or inside the box were excluded due to extremely low rates of occurrence in both experiments (Table S2). Each model included female behavior as the dependent variable with predator type, treatment order, and their interaction as fixed effects. Day of the year and male presence (yes or no) were included as covariates and female ID was included as a random effect to control for seasonal shifts in behavior, the presence of the female’s mate, and repeated sampling of the same individuals. We did not include location as a random effect in our models because all of our nest boxes are located in similar edge habitats and the sample size for each individual site is small (N<10), which substantially reduced our statistical power to interpret any potential differences. We applied an alpha value of 0.05 for all analyses.

The number of hits and flyovers as well as the closest approach to the decoy were not normally distributed owing to a large amount of zeros. Because transformations could not resolve this issue, we used GLMMs that assumed a Poisson distribution to analyze those behaviors. Closest approach scores were multiplied by 100 before the analysis to generate integers, which are required for the Poisson distribution. Time within 5m of the decoy was analyzed with LMMs that assumed a Gaussian distribution. Although the model residuals marginally failed a Shapiro-Wilk test of normality (package: “stats”), we visually inspected the residuals to confirm that they approximated a normal distribution. An abundance of evidence has indicated that LMMs are robust to violations of this assumption, except in cases where the sample sizes are prohibitively small or the distributions are bimodal, neither of which occurred here (Schielzeth et al. 2020; Knief and Forstmeier 2021; and further citations within). Time to return to the nest was also analyzed with LMMs that assumed a Gaussian distribution, but we square root transformed the data to achieve normality of the residuals based on Shapiro-Wilk tests and visual inspection of the residuals. The same transformation of the time within 5 m data made the residuals less normal, which is why we did not transform the data for that behavior. Because alarm calls were scored as a categorical variable (yes or no), we used GLMMs that assumed a binomial distribution.

To further evaluate model assumptions across our analyses, we tested for homoscedasticity using Levene’s test (package: “car”). Homoscedasticity was met for all models in experiment 1 but none of the models in experiment 2. These violations were seemingly due to the uniformly weak responses elicited by the hawk decoy in experiment 2. Although these violations likely had minimal effects on the validity of the models (Schielzeth et al. 2020; Knief and Forstmeier 2021), we ran a separate series of nonparametric Wilcoxon tests (package: “stats”) that lack the assumptions of normality and homoscedasticity to confirm the results of the mixed-effects models. For each behavior, we split the data based on treatment order and analyzed responses to each predator for the first trial separately from responses to the predators for the second trial. This approach allowed us to isolate the fixed effect of interest (predator type) from the fixed effect of order and the random effect of female identity. The qualitative results of these nonparametric tests were nearly identical to the mixed-effects models (Tables S5-6).

We also used nonparametric Wilcoxon tests to determine whether the number of neighboring conspecific and heterospecific individuals and the number of heterospecific species that responded to the trials differed between predator types in both years. This nonparametric test was chosen due to the lack of normality and low variance in these datasets caused by a large proportion of zero or near zero values. We considered applying a GLMM with a binomial distribution, but decided against this approach because it did not allow us to analyze both the total number of heterospecifics and the number of heterospecific species that responded because both reduced data sets (i.e., presence or absence of heterospecifics) would have been identical.

### Ethical Note

All procedures were reviewed and approved by the Ohio Wesleyan University Institutional Animal Care and Use Committee (Protocol Numbers: Spring_2020-21-01_Reichard and 05-2021-06), U.S. Federal Bird Banding Lab (Permit Number: 24035), and State of Ohio Department of Natural Resources (Permit Numbers: 23-009, 23-010). To minimize stress in the free-living birds used in this study, we attempted to capture and mark only the male of each pair, which was the minimum required to facilitate individual identification during our behavioral trials. All birds were immediately removed from mist nets after capture, processed rapidly to limit handling time, and then safely released at the capture site. No adverse effects were observed from these widely established techniques. Although our behavioral trials involved the observation of a simulated predator at the nest box, all subjects returned to normal parental care after the completion of the trial and no nests were abandoned as a result of these experiments.

## Results

### Experiment 1: Chipmunk v. Snake Decoy

Female wrens exhibited no significant differences in the number of hits (χ^2^_1_ =1.76, *P*=0.18), number of flyovers (χ^2^_1=2.34,_ , *P=*0.13), time spent within 5 meters of the nest box (F_1,39.1_=2.68, *P*=0.11), or in their closest approach ( ^2^ =2.46, *P*=0.12) to the snake and chipmunk decoys (Figure 1A-D). There was also no difference in the number of females (36 of 38 total for both predators; 94.7%) that produced alarm calls in response to each predator (χ^2^_1_<0.001, *P*>0.98). After the predator decoys were removed, we found no significant difference between predator types in the amount of time females took to re-enter the nest box (F_1,32.6_=0.68, *P*=0.42; Figure 2A).

**Figure 1.**
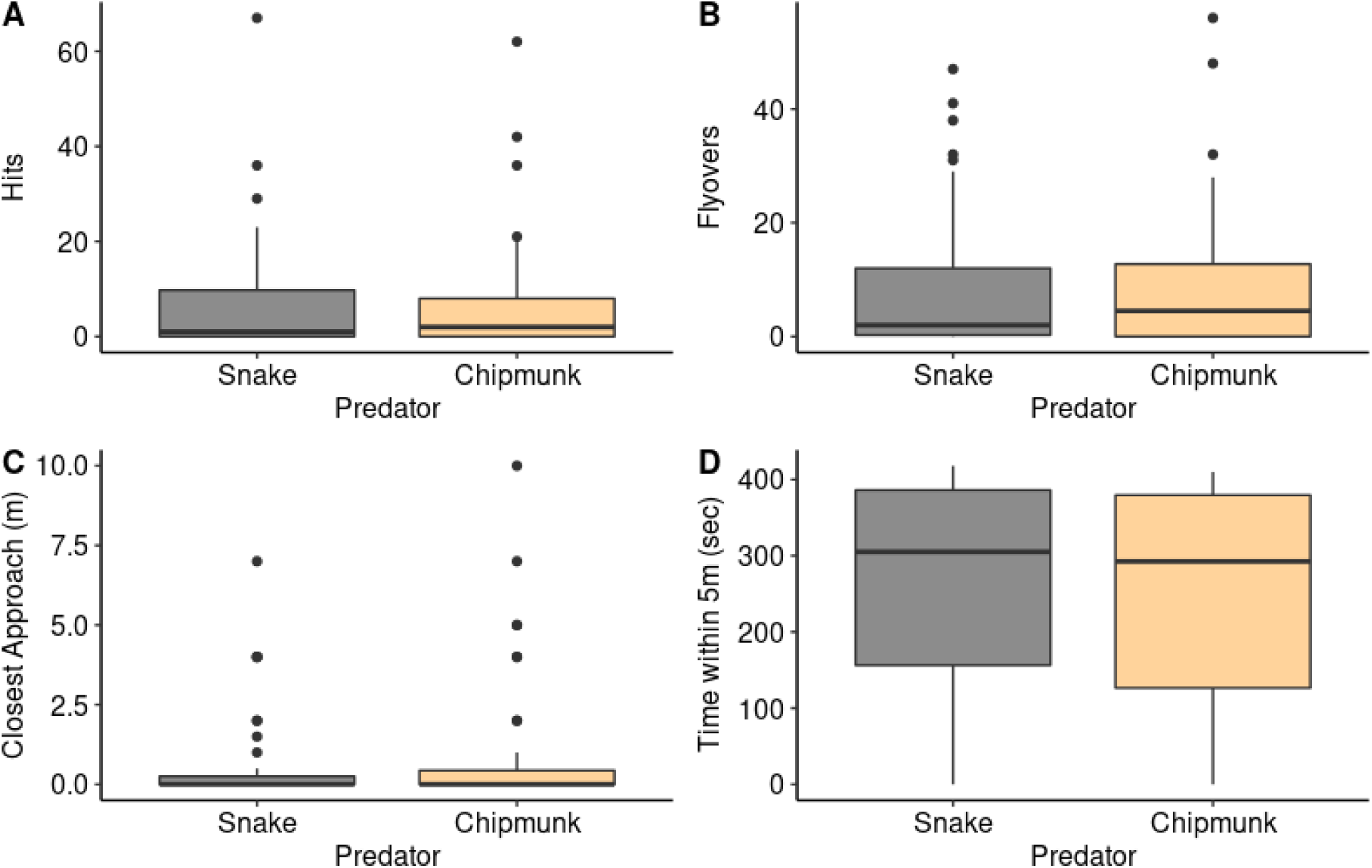
Responses of female house wrens to a snake or chipmunk decoy on top of their nest box during a seven-minute trial. There were no significant differences in the number of (A) hits (P=0.18), (B) flyovers (P=0.12), (C) how closely females approached each model (P=0.12), or (D) how much time females spent within 5 m of each model (P=0.11). The thick center line represents the median, and the box encloses the interquartile range. Whiskers show the range of data within 1.5 times the interquartile range, and dots are data points exceeding that range.

**Figure 2.**
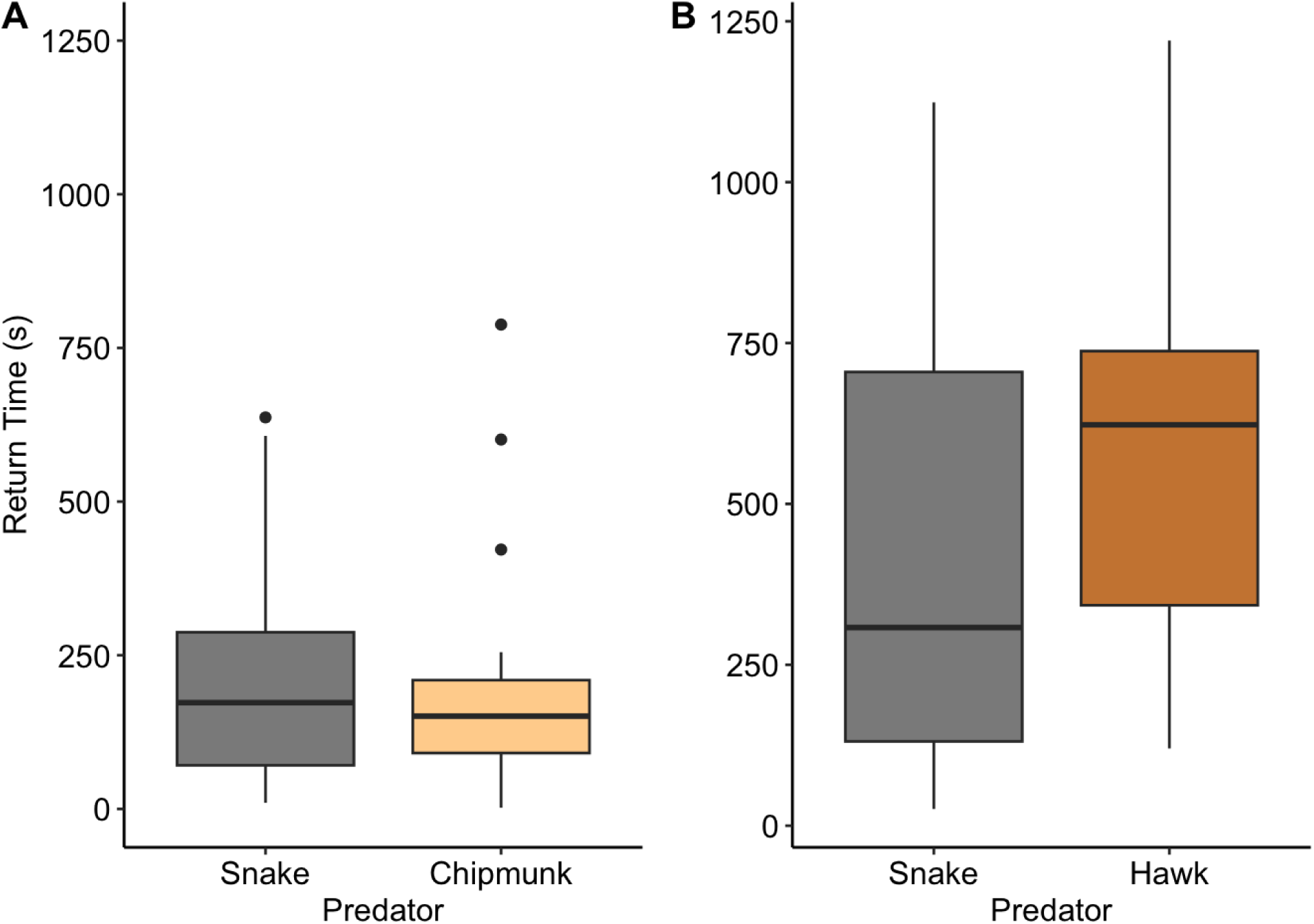
No detectable differences in how long it took female house wrens to re-enter the nest box after encountering a snake decoy versus a chipmunk decoy (P=0.42) or a hawk decoy (P=0.41). Box plots as in Fig. 1.

Day of the year had a significant effect on the number of hits ( ^2^ =8.42, *P*=0.003) with females hitting the decoy less often later in the breeding season. Similarly, females took longer to return to the nest box later in the season (F_1,27_=8.87, *P*=0.006). There was no detectable effect of date on any other behaviors (*P*≥0.46 in all cases; Table S3). The presence of the female’s mate had a significant effect on the time spent within 5 m of the decoy (F_1,34_=4.20, *P*=0.048) and how quickly the female returned to the nest ( ^2^ =7.86, *P*=0.009). When their mate was present, females spent more time within 5 m of the decoy and they returned to the nest faster. There was no detectable effect of male presence on any other behaviors (*P*≥0.13 in all cases; Table S3). For all six behaviors, we found no evidence for significant effects of treatment order (*P*≥0.055 in all cases; Table S3) or significant interactions between treatment order and predator type (*P*≥0.13 in all cases; Table S1). In one exception, we noted a marginal effect of treatment order on the closest approach to the decoy with females tending to approach more closely during the first trial (χ^2^_1_=3.67, *P*=0.055).

The number of conspecific (*W*=724.5, *P*=0.98) and heterospecific individuals (*W*=693.5, *P*=0.74) and the number of heterospecific species (W=700.5, *P*=0.80) that responded to the predator decoys did not differ detectably between the two predator types.

### Experiment 2: Hawk v. Snake Decoy

Similar to Experiment 1, female wrens frequently performed hits and flyovers while defending against the snake decoy, with 25 out of 41 females (61.0%) hitting the snake model at least once and 38 out of 41 females (92.7%) flying over at least once. Notably, zero of the female wrens performed hits or flyovers while defending against the hawk decoy. Our GLMM approach could not assess statistical significance with this data structure (i.e. no variation in response to the hawk), but our nonparametric analysis confirmed that females hit and flew over the snake significantly more often than the hawk (*P*<0.001 in both cases, Table S6; Figure 3A-B). Females approached significantly closer to the snake (χ^2^_1_ =21.8, *P* < 0.001; Figure 3C), but the amount of time spent within 5 meters was not significantly different from the response to the hawk (F_1,39.6_=1.28, *P*=0.27; Figure 3D). There was also no difference in the number of females (34 of 38 total for both predators; 89.5%) that produced alarm calls in response to each predator (χ^2^_1_ =0.10, *P*=0.75). After the removal of the predator decoys, female wrens did not differ detectably in the time to re-enter the nest box after the predator was removed (F_1,37.5_=0.70, *P*=0.41; Figure 2B). Finally, we found no evidence for an effect of date (*P*≥0.09 in all cases; Table S4), the presence of the female’s mate (*P*>0.27 in all cases; Table S4), treatment order (*P*>0.75 in all cases; Table S4) or a predator type by treatment order interaction (*P*>0.50 in all cases; Table S4) for any behaviors.

**Figure 3.**
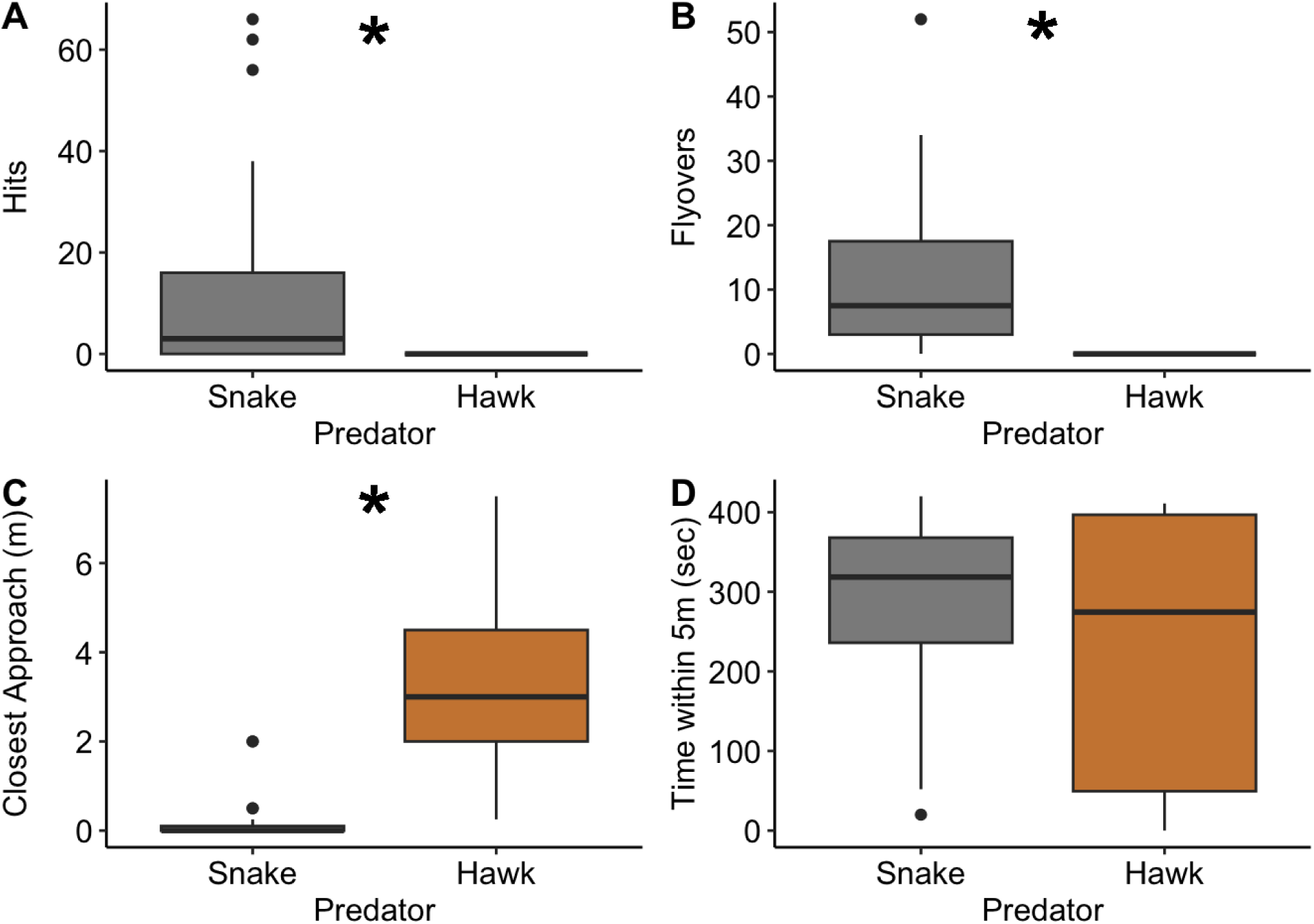
Responses of female house wrens to a snake or hawk decoy on top of their nest box during a seven-minute trial. Female house wrens performed significantly more (A) hits (P<0.001) and (B) flyovers (P<0.001) against the snake decoy than against the hawk decoy. Female wrens also (C) made significantly closer approaches to (P<0.001), but (D) they did not significantly differ in the time spent within 5 meters of the decoys (P=0.27). Box plots as in Fig. 1. *P<0.001.

Significantly more heterospecific individuals responded to the hawk decoy than to the snake decoy (*W*=531.5, *P*=0.04), but there was no detectable difference in the number of heterospecific species that responded to each predator (*W*=599.5, *P*=0.18). In contrast, significantly more conspecific individuals mobbed the snake decoy than mobbed the hawk (W=908, *P*=0.01).

## Discussion

Defending offspring from predators can be dangerous, and parents that accurately assess the risks posed by different predators and scale their responses appropriately should have higher lifetime fitness (Montgomerie and Weatherhead 1988; Clutton-Brock 1991; Caro 2005). We found evidence for risk assessment and plastic antipredator behavior in nesting house wrens. Female wrens responded with similar levels of aggressive nest defense behavior when encountering a simulated snake or chipmunk at their nest box, both of which posed a higher risk to offspring than adults. In contrast, the wrens responded significantly less aggressively to a simulated hawk that posed a higher risk to adults than nestlings. This plasticity in nest defense behavior is potentially adaptive because parents that avoid high risk predators like the hawk are more likely to live longer and have higher lifetime fitness despite the potential costs to the current clutch of offspring (Trivers 1974; Montgomerie and Weatherhead 1988).

Collectively, these results are consistent with the predictions of parental investment theory (Montgomerie and Weatherhead 1988; Caro 2005), and similar parental risk assessment in different contexts has been previously observed in house wrens and other species (Lima 2009; Mahr et al. 2015; Duré Ruiz et al. 2018).

We observed substantial individual variation among females in how strongly they responded to each of the simulated predators. For example, the most aggressive females hit the snake and chipmunk decoys over 50 times in a seven-minute period while the least aggressive females never dove at or hit the decoys. Some of this variation could be explained by parental investment theory, which predicts a positive relationship between the intensity of nest defense and various life history traits such as clutch size, nestling age, and adult age and longevity (Montgomerie and Weatherhead 1988, Caro 2005). However, testing these predictions was not a main focus of our study; rather, we sought to control for differences in parental investment across the nestling stage. As a result, we only captured a fraction of the possible variation in clutch size (80% of pairs defended 4-6 offspring) and nestling age (85% of offspring were 3-6 days post hatch), which limits our ability to draw conclusions about the relationship between aggression and current reproductive investment. In Experiment 1, we observed that females hit the predator decoy less and took longer to return to the nest when trials occurred later in the breeding season. This result is consistent with a redirection of investment away from offspring towards the individual, which could be adaptive in a species with multiple breeding attempts. House wrens, however, are short-lived with an average lifespan of less than two years (Johnson 2020), so the benefit, if any, of this shift remains unclear. Alternatively, reduced effort later in the season could result from a seasonal decline in female condition due the cumulative effects of multiple breeding attempts (Laiolo et al. 2004, Mitchell et al. 2012). More research is necessary to evaluate how these energetic tradeoffs might shape the different components of antipredator behavior, and in turn, affect both adult and offspring survival in house wrens and other species (Lind and Cresswell 2005).

Two important exceptions to the persistent variation among individuals in nest defense behavior occurred with (1) the production of alarm calls when any predator decoy was present, and (2) the behavioral responses to the simulated hawk. In both experiments, females almost universally produced alarm calls despite varying levels of risk posed by the different predators to the adults and their nestlings. Alarm calls are a lower risk behavior than the other nest defense behaviors sampled here, which might explain their consistent production. When produced near the nest, alarm calls can serve many adaptive functions including distracting the predator (Fasanella and Fernández 2009), inducing neighboring birds to mob the predator (Templeton et al. 2005), and alerting the young inside the nest that a predator is nearby (Suzuki 2011). Previous research in another house wren population found that parents produced alarm calls even in the presence of a non-predatory heterospecific bird at their nest (Fasanella and Fernández 2009), suggesting that alarm calls might function as a general response to any disturbance at the nest box. However, wren parents also produced more alarm calls in response to a potential predator rather than a non-predatory control bird (Fasanella and Fernández 2009) and more alarm calls were produced as a simulated snake moved closer to the nest (Fernández and Carro 2022). Although we did not measure whether the quantity of alarm calls differed between predator types in our study, it seems clear that alarm calls are a consistent and important component of the antipredator response, and the intensity of these calls might signal information about the predator type and level of risk posed to the nestlings (Fasanella and Fernández 2009, Fernández and Carro 2022, Fernández et al. 2023).

When responding to the hawk decoy, females varied in how closely they approached and how much time they spent within five meters of the hawk, but none of the 41 females ever hit the hawk or even flew over it. This consistency among individuals indicates that when the predatory threat surpasses a certain level it is possible to see uniformity in a typically plastic behavior. Furthermore, the total absence of physical aggression towards the hawk supports the hypothesis that selection favors risk assessment, likely by acting against high-risk behaviors, such as closely approaching or physically attacking a dangerous predator (Desrocher et al. 2002; Mahr et al. 2015). However, our observations of the community-level response to the hawk and other simulated predators show that risk assessment is not uniform across songbird species.

Neighboring conspecifics and heterospecific songbirds frequently responded to our simulated predators by approaching and alarm calling, but very few approached closely enough to attack or fly directly over the decoys, including the snake and chipmunk (RCE, DGR, JHF, personal observations). This observation is consistent with mobbing behavior, and the reliance on alarm calls rather than overt aggression likely reflects the reduced threat posed to these neighbors by a predator that was not near their own offspring (Colombelli-Négrel et al. 2010; Kryštofková et al. 2011). Similar to the antipredator behavior of the wren parents, the number of conspecifics and heterospecifics that responded did not differ significantly between the snake and chipmunk. In contrast, more conspecifics responded to the snake while more heterospecifics responded to the hawk in our second experiment. This result mirrors the cautious behavior of house wrens when responding to a hawk at their nest box, which likely explains the lack of mobbing behavior from conspecific neighbors. In contrast, larger heterospecific songbirds including tree swallows (*Tachycineta bicolor*), northern cardinals (*Cardinalis cardinalis*), and blue jays (*Cyanocitta cristata*) appeared in higher numbers when the hawk was present, and tree swallows frequently dove at the hawk, which was in stark contrast with all other species (RCE, DGR, JHF, personal observations).

The heterospecific response to the hawk was surprising given that Cooper’s hawks primarily forage for larger songbirds (Roth and Lima 2003; Rosenfield et al. 2020), which elevated the risk of predation for those that approached. Although some heterospecifics were likely defending nearby nests from the hawk (e.g., tree swallows from nearby boxes with active nests), others may have been gathering information through eavesdropping behavior, which can be used to inform future decisions related to predation threats (Magrath et al. 2015). House wrens, as an insectivorous species that must be highly vigilant in search of prey, may be better able to detect predators than less vigilant species or those that spend more time in denser substrate, thereby providing more accurate spatial information for eavesdropping heterospecifics (Magrath et al. 2015; Johnson, 2020). Future experiments could examine the ecological relationship between defensive cooperation, social learning, and information flow through social networks of different avian communities.

Reactive nest defense behavior can be subdivided into (1) behaviors that distract or redirect the predator towards the signaler and (2) behaviors that involve acts of aggression towards the predator (Caro 2005; Lima 2009). Smaller organisms like house wrens may rely on distraction rather than attacking predators above a certain size because they lack the body size or armaments necessary to successfully drive the predator away (Montgomerie and Weatherhead 1988; Winkler 1992). If the likelihood of physically deterring the predator is low, then any amount of risk to the parent from attacking the predator could justify a distraction-based antipredator response or even no response at all. However, it is unlikely that size thresholds universally govern antipredator behavior given that certain species of limited size and armaments frequently attack larger predators, including humans (Levey et al. 2009, Lawson et al. 2021). In addition to size differences, birds of prey are substantially more mobile in the air than terrestrial predators, which further elevates the risks associated with physical aggression and could explain distraction-based responses regardless of an avian predator’s size (Mahr et al. 2015). More broadly, risk-taking thresholds and antipredator behavior can vary according to other factors such as predator abundance, resource availability, or competition as adult birds employ strategies that balance the reproduction-survival tradeoff (Dı az et al., 2013; Mikula et al., 2018). Future work should continue to focus on elucidating the proximate and ultimate causes of persistent intra- and interspecific variation in parental antipredator behavior and risk assessment.

**Table 1.** Mean ± standard deviation (range) for the number of conspecific and heterospecific individuals and heterospecific species that responded to the predator stimuli in Experiment 1.

| Observation | Snake | Chipmunk |
| --- | --- | --- |
| Conspecific Individuals | 0.53 ± 1.0<br>(0-5) | 0.47 ± 0.8<br>(0-3) |
| Heterospecific Individuals | 1.08 ± 2.7<br>(0-15) | 0.76 ± 1.4<br>(0-6) |
| Heterospecific Species | 0.61 ± 1.0<br>(0-5) | 0.58 ± 0.9<br>(0-4) |

**Table 2.**
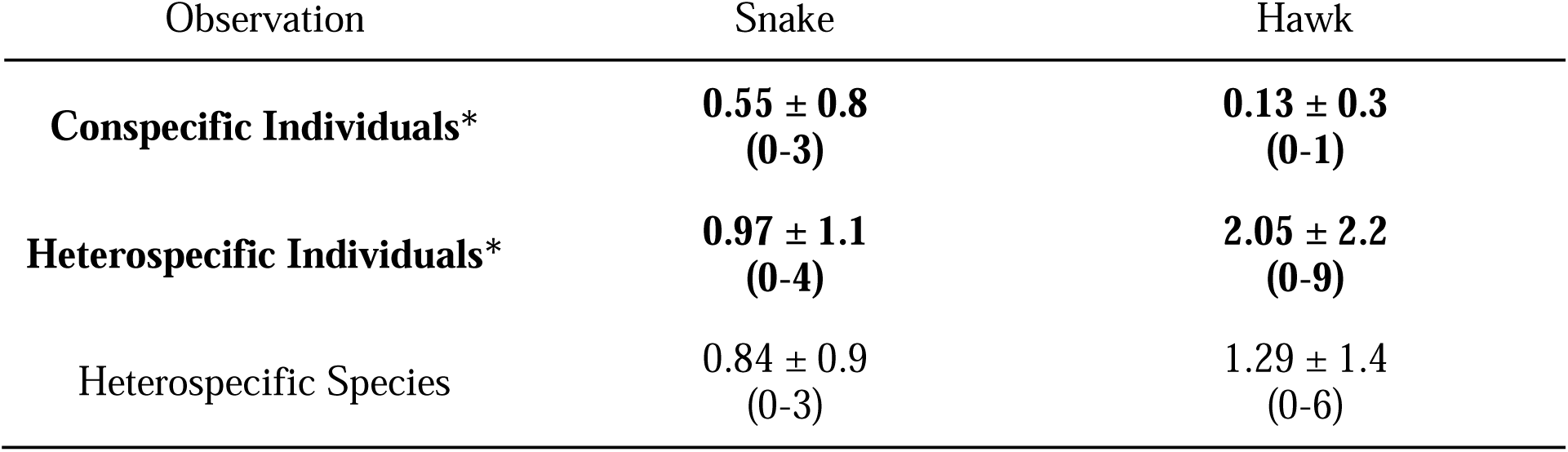
Mean ± standard deviation (range) for the number of conspecific and heterospecific individuals and heterospecific species that responded to the predator stimuli in Experiment 2. Bold text denotes *P*<0.05.

| Observation | Snake | Hawk |
| --- | --- | --- |
| **Conspecific Individuals\*** | **$0.55 \pm 0.8$**<br>**(0-3)** | **$0.13 \pm 0.3$**<br>**(0-1)** |
| **Heterospecific Individuals\*** | **$0.97 \pm 1.1$**<br>**(0-4)** | **$2.05 \pm 2.2$**<br>**(0-9)** |
| Heterospecific Species | $0.84 \pm 0.9$<br>(0-3) | $1.29 \pm 1.4$<br>(0-6) |

## Data Availability Statement

The data collected and analyzed to support these findings are freely available in the Mendeley Data Repository at DOI:10.17632/zjb5ynz5n4.2.

## Acknowledgements

We thank Ohio Wesleyan University, the Big Walnut School System, and the Davis, Koban, and Fink families for access to facilities and field sites. We thank Holly Keating for analyzing videos, Tamaki Yuri for preparing the taxidermied Cooper’s Hawk, Lily Bonar for field assistance, Chandler Carr for measuring our nest boxes, and Eric Gangloff for advice on statistics. This project was funded by the Ohio Wesleyan University Summer Science Research Program.

## Ethics Approval Statement

All procedures were reviewed and approved by the Ohio Wesleyan University Institutional Animal Care and Use Committee (Protocol Numbers: Spring_2020-21-01_Reichard and 05-2021-06), U.S. Federal Bird Banding Lab (Permit Number: 24035), and State of Ohio Department of Natural Resources (Permit Numbers: 23-009, 23-010).

## Supplementary Materials

### Statement on STRANGE framework

Studies of animal behavior can be misinterpreted due to sampling biases, and the STRANGE framework provides a method for identifying these potential issues (Webster and Rutz 2020). All subjects in this study were free-living songbirds, and we included all individuals that nested on our study sites regardless of age or nesting location, making it unlikely that social background biased our results. As detailed above, a small number of individuals were excluded, namely those that failed to respond within 7 minutes during Experiment 1 before our protocol was adjusted to be more inclusive or those that responded so forcefully that they knocked the predator decoy from the box. Thus, we may have failed to sample a subset of both low and high responders. Although this approach may have biased our sample towards intermediate responders, we think this potential bias is unlikely to have skewed our results for a few reasons. First, our sample size is large for a field study and it includes individuals that responded both aggressively and weakly to the predator decoys, so it is unlikely that we failed to sample the extremes of behavioral variation in this population.

Second, we conducted repeated-measures sampling, which required subjects to be excluded from all treatments rather than only those trials where we were unable to collect data. This approach limits the likelihood of bias as each individual effectively serves as its own control. With respect to seasonal changes in behavior, we sampled evenly across the entire breeding season and included measurement date as a covariate in our models to test for this effect.

Because each subject experienced two sampling events it is possible that habituation affected responses to the second treatment. We limited this effect by presenting trials on separate days in a counterbalanced order and included treatment order in our models to test for an effect of habituation.

It is also possible that carryover effects caused by sampling the same individuals between years could bias the behavioral responses in Experiment 2. We think that this outcome is unlikely given the length of time between the two experiments, but if the same females were sampled in both years carryover effects remain possible. It is difficult to determine how many females were sampled in both experiments because only 13 of 47 females (27.7%) were banded in experiment one, including six females banded in previous breeding seasons. Of those banded females, only one (7.7.% return rate) was definitely sampled in experiment two. It is likely that some of the unbanded females also returned to be sampled again, and some inferences can be drawn from males, which were banded at much higher rates. In experiment one, 43 of 47 males (91.5%) were banded and eight males returned to be sampled again in experiment two (18.5% return rate). Based on these data, it seems likely that 10-20% of females were sampled in both experiments.

**Table S1.** Locations of field sites and number of nest boxes present.

| Site | Location | Number of Boxes |
| --- | --- | --- |
| Big Walnut Elementary School, Sunbury, Ohio | 40°14'02.6"N<br>82°51'53.1"W | 15 |
| Big Walnut High School, Sunbury, Ohio | 40°14'00.9"N<br>82°51'37.3"W | 41 |
| Davis Family Property, Sunbury, Ohio | 40°15'50.2"N<br>82°49'52.8"W | 36 |
| Fink Family Property, Marysville, Ohio | 40°16'42.4"N<br>83°15'43.3"W | 21 |
| Kraus Research Preserve, Delaware, Ohio | 40°11'45.8"N<br>83°03'47.1"W | 42 |
| Ohio Wesleyan University and nearby homes, Delaware, Ohio | 40°17'47.7"N<br>83°03'53.8"W | 25 |

**Table S2.** Summary of the number of trials that include an observation of time spent in or on the next box. Experiment 1 tested responses to a simulated snake and chipmunk. Experiment 2 tested responses to a simulated snake and hawk.

| Experiment | Time spent in the nest box | Time spent on the nest box |
| --- | --- | --- |
| 1 | 4/76 (5.3%) | 12/76 (15.8%)<sup>1</sup> |
| 2 | 4/76 (5.3%) | 10/76 (13.2%)<sup>2</sup> |

<sup>1</sup>9 of 12 observations were for 10 seconds or less (75%) <sup>2</sup>7 of 10 observations were for 10 seconds or less (70%)

**Table S3.** Analysis of model fixed effects and covariates for Experiment 1. *P<0.05.

| Behavior | Day of the Year |  | Male Presence |  | Order |  | Interaction with Order |  |
| --- | --- | --- | --- | --- | --- | --- | --- | --- |
|  | X<sup>2</sup> or F | P | X<sup>2</sup> or F | P | X<sup>2</sup> or F | P | X<sup>2</sup> or F | P |
| Hits | 8.42 | 0.004* | 0.38 | 0.54 | 2.11 | 0.15 | 1.87 | 0.17 |
| Flyovers | 0.02 | 0.88 | 2.09 | 0.15 | 2.11 | 0.15 | 2.34 | 0.13 |
| Time within 5 m | 0.33 | 0.57 | 4.20 | 0.048* | 0.004 | 0.95 | 2.30 | 0.14 |
| Closest Approach | 0.55 | 0.46 | 2.25 | 0.13 | 3.67 | 0.055 | 1.91 | 0.17 |
| Latency to Return | 8.87 | 0.006* | 7.86 | 0.009* | 1.59 | 0.23 | 0.67 | 0.42 |
| Alarm Calls | 0.18 | 0.67 | 1.01 | 0.32 | 0.10 | 0.76 | <0.001 | 0.99 |

**Table S4.** Analysis of model fixed effects and covariates for Experiment 2.

| Behavior | Day of the Year |  | Male Presence |  | Order |  | Interaction with Order |  |
| --- | --- | --- | --- | --- | --- | --- | --- | --- |
|  | X<sup>2</sup> or F | P | X<sup>2</sup> or F | P | X<sup>2</sup> or F | P | X<sup>2</sup> or F | P |
| Time within 5 m | 2.16 | 0.15 | 0.54 | 0.47 | 0.02 | 0.89 | 0.47 | 0.50 |
| Closest Approach | 2.83 | 0.09 | 1.22 | 0.27 | 0.006 | 0.94 | 0.22 | 0.64 |
| Latency to Return | 1.06 | 0.31 | 0.10 | 0.75 | <0.001 | 0.98 | 0.001 | 0.97 |
| Alarm Calls | 0.13 | 0.71 | 0.07 | 0.80 | 0.10 | 0.75 | 0.11 | 0.74 |

**Table S5.** Results of nonparametric analysis of Experiment 1 behavior.

| Behavior | First Trial |  | Second Trial |  |
| --- | --- | --- | --- | --- |
|  | W | P | W | P |
| Hits | 200 | 0.41 | 118 | 0.09 |
| Flyovers | 220.5 | 0.15 | 126.5 | 0.17 |
| Time within 5 m | 204 | 0.35 | 168 | 0.91 |
| Closest Approach | 147 | 0.39 | 210.5 | 0.23 |
| Latency to Return | 116 | 0.82 | 109 | 0.98 |

**Table S6.** Results of nonparametric analysis of Experiment 2 behavior. *P<0.05.

| Behavior | First Trial |  | Second Trial |  |
| --- | --- | --- | --- | --- |
|  | W | P | W | P |
| Hits | 280 | <0.001* | 288 | <0.001* |
| Flyovers | 350 | <0.001* | 351 | <0.001* |
| Time within 5 m | 223 | 0.21 | 170 | 0.78 |
| Closest Approach | 4 | <0.001* | 8.5 | <0.001* |
| Latency to Return | 92 | 0.048* | 111.5 | 0.19 |

